# Engineering Alignment has Mixed Effects on hiPSC-CM Maturation

**DOI:** 10.1101/2022.09.26.509194

**Authors:** Nikhith G. Kalkunte, Talia E Delabmbre, Sogu Sohn, Madison Pickett, Sapun Parekh, Janet Zoldan

## Abstract

The potential of human induced pluripotent stem cell differentiated cardiomyocytes (hiPSC-CMs) is greatly limited by their functional immaturity. Strong relationships exist between CM structure and function, leading many in the field to seek ways to mature hiPSC-CMs by culturing on biomimetic substrates, specifically those that promote alignment. However, these *in vitro* models have so far failed to replicate the alignment that occurs during cardiac differentiation. We show that engineered alignment, incorporated before and during cardiac differentiation, affects hiPSC-CM electrochemical coupling and mitochondrial morphology. We successfully engineer alignment in differentiating hiPSCs as early as Day 4. We uniquely apply optical redox imaging to monitor the metabolic changes occurring during cardiac differentiation. We couple this modality with cardiac-specific markers, which allows us to assess cardiac metabolism in heterogeneous cell populations. The engineered alignment drives hiPSC-CM differentiation towards the ventricular compact CM subtype and improves electrochemical coupling in the short term. Moreover, we observe glycolysis to oxidative phosphorylation switch throughout differentiation and CM development. On the subcellular scale, we note changes in mitochondrial morphology in the long term. Our results demonstrate that cellular alignment accelerates hiPSC-CM maturity and emphasizes the interrelation of structure and function in cardiac development. We anticipate that combining engineered alignment with additional maturation strategies will result in improved development of mature CMs from hiPSC and strongly improve cardiac tissue engineering.

**Impact Statement:** This work demonstrates the mixed effect of engineered structure in inducing matured function of human induced pluripotent stem cell -derived cardiomyocytes. Isolating the impacts of hiPSC-CM alignment on functionality is a necessary step in optimizing the culture conditions to develop cardiac cell therapies. Furthermore, our work has broader implications concerning how we understand the impact of mechanical microenvironments on stem cell differentiation and development.

## Introduction

Cardiovascular disease (CVD) is the leading cause of death worldwide and affects over 80 million Americans yearly.^1^ The impact of CVD is magnified by the heart’s inability to repair and self-regenerate, resulting in permanent cardiac output deficiencies.^2^

Human induced pluripotent stem cells (hiPSCs) look to be possible solutions to CVD by providing a source of non-immunogenic cells that can differentiate into various cells found in the heart. Yet, the functional immaturity of hiPSC differentiated cardiomyocytes (hiPSC-CMs) has stymied their translation to the clinic. iPSC-CMs exhibit various functional deficiencies as compared to adult CMs.^3,4^ More specifically, poor electrochemical coupling in hiPSC-CMs can be tied to the lack of t-tubule networks,^4,5^ slower calcium transients,^6^ and low expression of sarco/endoplasmic reticulum Ca2+-ATPase 2a (SERaCA2a).^7^ hiPSC-CMs are also metabolically immature, exhibiting lower mitochondrial content and relying on the less efficient glycolysis rather than fatty acid oxidation for ATP generation.^8^

Improper CM alignment has been theorized to be one of the sources of these functional deficiencies. The relationship between the anisotropic alignment of CMs and their functionality has been extensively studied.^3,7,9–11^ Protein machinery that supports electrochemical coupling is directly tied to cell alignment. Gap junctions are concentrated at longitudinal ends of the CMs, facilitating unidirectional signal propagation.^12^ Indeed, engineering alignment in isolated primary CMs has improved electrochemical coupling and calcium handling.^13,14^ Similarly, metabolism in CMs shows interesting mechanoresponsive characteristics. Mitochondria within CMs are collocated and aligned with sarcomeres and the sarcoplasmic reticulum to promote efficient calcium release and contraction.^15^

Importantly, cellular alignment occurs progressively throughout cardiogenesis. Murine models showcase the alignment of CMs in the mouse heart starting as early as embryonic day 12.5 and lasting through postnatal day 4.^16^ This suggests similar alignment likely occurs in human heart development. Thus, *in vitro* efforts to clarify the impact of alignment on human CM functionality should also promote CM alignment during differentiation rather than post CM differentiation, as most studies have done. Indeed, the differentiation of hiPSC-derived cardiac progenitor cells (hiPSCs-CPs) on aligned fibrous scaffolds improved calcium handling compared to unaligned controls.^17^ Similarly, the metabolism of stem cells is dynamic over differentiation.^18^ The CM mitochondrial network re organizes over development, transitioning from a random sparse mitochondria to an ordered, structured network of mitochondria.^19^

We therefore hypothesized that introducing alignment during hiPSC differentiation into CMs will impact their function and metabolism. Assessing hiPSC-CM maturation over differentiation requires precise measurement techniques to differentiate CM behavior among other cell types. Electrochemical coupling assessment is easily facilitated by calcium-sensitive imaging.^20^ Measuring CM metabolism in heterogeneous cell culture is not possible using common ensemble metabolism measurements like oxygen consumption rate or glucose consumption assays. Instead, a CM-specific metabolic measurement tool is needed. Optical redox imaging (ORI) uses two-photon microscopy and the endogenous fluorescence of metabolic intermediates Nicotinamide adenine dinucleotide (NADH) and Flavin adenine dinucleotide (FAD+) to assess metabolic pathways. As glycolysis results in the reduction of NADH and FAD+ is synthesized in the electron transport chain, reporting the optical redox ratio (FAD+/NADH+FAD+) captures the metabolic preference of cells from 0 (glycolysis) to 1 (oxidative phosphorylation). The optical redox ratio (ORR) has been extensively used in cancer biology,^21–23^ and is strongly correlated with the concentrations of NAD+/NADH.^18^ Here, we combine cardiac-specific maker staining with ORR to assess changes in CM metabolism in heterogeneous cell populations.

Herein, we uniquely apply calcium imaging and ORI to investigate how alignment, incorporated prior to differentiation, affects hiPSCs-CM development and functionality. We show that alignment impacts hiPSC-CM subtype development and expose a temporal relationship between hiPSC-CM alignment and calcium handling. We further demonstrate no impact of alignment on hiPSC-CM metabolism but reveal impacts of alignment on mitochondrial morphology.

## Methods

### PCL Fiber Synthesis and Characterization

Random and aligned fiber substrates were created by electrospinning poly(caprolactone) (PCL) (Sigma Aldrich) onto a rotating collector mandrel. Electrospun fiber orientation and diameter were calculated using ImageJ/Fiji and MATLAB (Mathworks) to process scanning electron microscope (SEM) images as we described previously.^24^ Tensile testing was conducted via force ramping. Mounted samples were loaded into a UniVert mechanical tester (CellScale) with a 10N load cell. In tension mode, samples were stretched at 8N in a ramp fashion. Displacement and normal force were recorded over a stretch period of 30s. Stress and strain were calculated and plotted to calculate Young’s Modulus of samples. Aligned fibers were tested parallel and perpendicular to the direction of fiber alignment.

### Cell Seeding and Differentiation

Pre-sterilized and Matrigel-coated fibers were seeded with 125,000 hiPSCs per cm^2^. Tissue culture polystyrene (TCP) served as control substrates. HiPSCs were differentiated into CMs via Wnt modulation. ^25^ Briefly, hiPSCs were initiated with RPMI 1640 media supplemented with B27 no insulin (RB-) and CHIR99021 (12mM). Twenty-four hours after induction with CHIR, cells were transitioned to RB-media for forty-eight hours. Cells were then transferred to RB-media supplemented with 5mM iWP2 (inhibitor of WNT pathway) for forty-eight hours. Next, cells recovered in RB- for another forty-eight hours. Thereafter cells were cultured in RPMI supplemented with B27 and insulin (RB+).

### Immunostaining

Cells differentiated on fibers were fixed and stained on days 0,1,4,14, and 28 to assess differentiation progress, alignment, and functionality (Supplementary Table S1). Stained samples were imaged using an EVOS XL Core Cell Imaging System, Leica DMI6000 B, or Olympus FV3000 Confocal Laser Scanning Microscope.

### Quantitative reverse transcription–polymerase chain reaction

Total RNA was extracted from cells on fibers on days 14 and 28 of CM differentiation using a RNeasy Mini Kit (Qiagen), according to manufacturer’s protocol. Reverse transcription was performed using a High-Capacity cDNA Reverse Transcription Kit (Applied Biosystems). The MasterMix was made according to the manufacturer’s protocol. Quantitative PCR was performed using PrimeTime qPCR Primers (Integrated DNA Technologies) and PowerUp SYBR Green (Thermo Fisher) on a StepOnePlus Real-Time PCR System (Applied Biosystems). TATA-binding protein (TBP) was used as the endogenous control, and mRNA isolated from CMs differentiated on tissue culture polystyrene served as controls for differentiation on fibers. as the control reference sample. The complete list of primers used is available in Supplementary Table S2.

### Cell Alignment

To quantify cell alignment, cells seeded on random and aligned fibers were fixed and immunostained with rhodamine-phalloidin and Draq5. 20X images were collected on an Olympus FV3000 confocal microscope. Images were analyzed in Fiji/ImageJ and MATLAB using the Orientation J plug-in to determine the cell orientation index, as described previously.^24^

### Intracellular Calcium Transients Imaging

CMV-GCaMP2 transfected hiPSCs, a gift from Dr. Bruce Konklin, were used to visualize spontaneous intracellular calcium transients. Spontaneous CM beating was confirmed on all samples by GFP signal on day 10. Image sequences at 20 frames per second were captured using a Leica DMI6000 B microscope and a 10X Leica objective.

Image sequences were processed in MATLAB (Mathworks) using custom code to determine synchronicity, beating rate, calcium cycle duration, and decay constant. Synchronicity is calculated as the median absolute deviation (MAD) of the time of peak arrival at each beating pixel. Pixels with little calcium flux (low changes in fluorescence intensity over time) were filtered out as non-beating. Five time points per image sequence were analyzed. The beating rate (presented at beats per minute, BPM) was calculated by dividing the number of calcium peaks in each image sequence by the time length of each image sequence. Calcium cycle duration was calculated as the full-width half-max of each calcium peak. The decay constant was calculated by fitting the decreasing GFP signal after each peak to an exponential curve (*Ae^-t/t^*), where τ is the decay constant.

### Optical Redox Ratio

Metabolism was measured by evaluating the optical redox ratio (ORR) of cells via two-photon excitation microscopy. A home-built two-photon microscope leveraging an Olympus FV3000 confocal microscope, a tunable femtosecond laser (Insight X3 Dual, Spectra-Physics), and a UPlanSApo 20 X/ 0.75NA Olympus objective was used to collect endogenous metabolic intermediates. Briefly, the autofluorescence of metabolic intermediates NADH and FAD were measured by exciting at 750nm and 860nm, respectively. Emissions were split with a dichroic mirror collected at 460nm±60 (NADH) and 530nm±43 (FAD) filters, and the ORR was calculated as FAD/(NADH+FAD) on a pixel-by-pixel basis in the field of view. This value, ranging from 0 to 1, is an ensemble measurement related to cellular metabolism where an ORR~0 indicates metabolism via glycolysis, whereas an ORR~1 indicates metabolism via oxidative phosphorylation. CMs in the ORR field of view were uniquely identified by staining with CM surface maker SIRPa (1:100, Biolegend, 323802). We performed control studies to ensure that the SIRPa signal did not interfere with ORR measurement (Supplementary Fig. S4a). CM-specific ORR was calculated by binarizing the SIRPa signal, which was used as a mask to multiply the ORR signal pixel-wise. The average ORR signal of CMs differentiated on glass, random, and aligned fibers was calculated using a custom MATLAB code on days 4, 14, and 28. ORR sensitivity was confirmed by collecting ORR after dosing cells with metabolic modulators Oligomycin A (3μM) and Carbonyl cyanide-p-trifluoromethoxyphenylhydrazone (FCCP) (2μM). Cells were incubated in drug solutions for 30 minutes and analyzed as described above.

### Mitochondrial Analysis

The mitochondrial area was assessed by staining samples with MitoTracker™ Deep Red FM (Invitrogen) and imaging with confocal microscopy. Day 14 and Day 28, differentiated CMs were incubated with MitoTracker™ Deep Red FM (Invitrogen; 500nM) for 45 min at 37C. Samples were washed 3 times in sterile DPBS and stained with CM surface marker SIRPa. Importantly, cells were not fixed for SIRPa staining. SIRPa and MitoTracker images were collected using an Olympus confocal microscope and a LUMPLFLN 40XW/0.80 NA Olympus objective. The cardiac-specific mitochondrial area was found by processing collected images in Fiji/ImageJ. As for ORR images, MitoTracker images were masked by SIRPa images to select mitochondria within CMs only. Fiji was then used, coupled with MATLAB, to calculate the area of identified CM-specific mitochondria and the average mitochondrial area for each condition.

### Statistical Analysis

Statistical analysis was performed using unpaired two-sample t-tests in Prism 9 (GraphPad). Relationships with p<0.05 were considered significant. Data are present as mean ± standard error.

## Experiment

### Electrospun Fiber Characterization

Aligned and random electrospun fibers were analyzed for fiber orientation, diameter, and stiffness. Orientations analysis of fibers yielded significant differences between aligned (S = 0.74± 0.02) and random (S = 0.20±0.05) fibers. Fiber diameter varied significantly between aligned (0.69± 0.02μm) and random (1.18± 0.02μm) fiber. Finally, the Young’s modulus of fiber mats was significantly different between aligned (54.82± 9.97 MPa) and random (24.50± 2.25 MPa) fibers (Supplementary Fig. S1).

### Cardiac Differentiation

hiPSCs seeded onto electrospun fibers were differentiated into CMs via Wnt pathway modulation. Immunostaining on days 1, 4, and 14 shows the differentiation progression through mesoderm and committed cardiac phases. Day 1 samples stained for mesoderm marker Brachyury T show collocation of Brachyury T and nuclei. Day 4 staining of Mesoderm Posterior BHLH Transcription Factor 1 (MESP1) also showed translocation of the transcription factor into the nuclei (Fig. 1A). Spontaneous beating, as indicated by GFP fluorescence, was seen in all analyzed samples on Day 10. Analysis of Day 14 CMs shows high gene and protein expression of the canonical cardiac marker, cardiac troponin T (cTnT). Immunostaining for cTnT shows its prevalence in both random and aligned samples (Fig. 1A). Immunostaining for cell markers vimentin, CD31, and α-smooth muscle actin were also positive in both ransom and aligned samples, indicating a heterogeneous cell population (Supplementary Fig S2). Gene expression analysis shows a significant threefold increase in expression of *TNNT2* in cells differentiated on random (3.73±0.50) or aligned (3.38±0.30) fibers compared to control (Fig. 1B). Cardiac subtype analysis of Day 14 CMs reveals that fiber alignment impacts specialization of CMs into ventricular cardiac tissues. Expression of ventricular cardiac marker *IRX4* is upregulated twelvefold in CMs differentiated on aligned fibers and fivefold in CMs differentiated on random fibers, relative to control. Similarly, atrial cardiac marker *NR2F2* is downregulated fivefold in CMs differentiated on aligned fibers, and threefold in CMs differentiated on random fibers, relative to control (Fig. 1B). Alignment further impacts the specialization of ventricular CMs into compact ventricular tissue. Expression of trabecular cardiac tissue marker *NNPA* is downregulated sixfold in CMs differentiated on aligned fibers, while only threefold in CMs differentiated on random fibers. Finally, both CMs differentiated on aligned (twofold) and random (twofold) fibers exhibited upregulation in compact cardiac tissue marker *HEY2* relative to control.

**FIG. 1.**
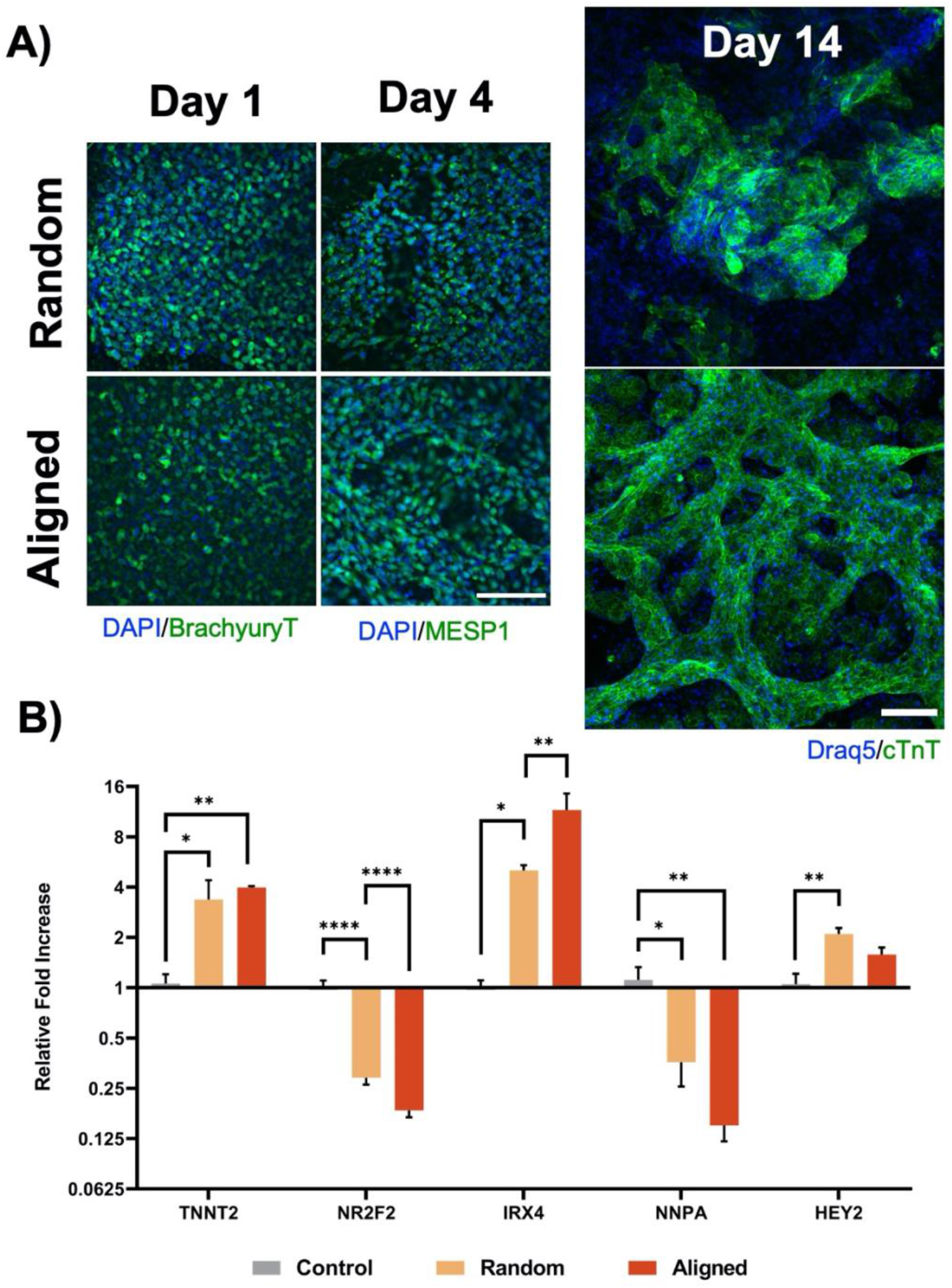
hiPSC are differentiated into cardiomyocytes on electrospun fibers and are assessed for subtype. **(A)** Representative images of hiPSC at Day 1, 4 and 14 of differentiation. Immunostaining for Brachyury T and MESP1 confirm progression of cardiac differentiation. Immunostaining cTnT proves successful cardiac differentiation on aligned and random electrospun fibers (scale bar = 100μm). **(B)** Effect of cell alignment at day 14 on mRNA expression of cardiac subtype genes (n=6). *p<0.05,**p<0.01,****p<0.0001.

### Cell Alignment

To evaluate the impact of substrate anisotropy on cell alignment, cells were immunostained and imaged for actin filaments at Day 0, 4, 14, and 28 of differentiation (Fig. 2A). These images show that cell alignment increases over time, but this trend is promoted or attenuated based on substrate alignment. As expected, at day 0, no significant differences in alignment are seen. By day 4, cells start to take on the alignment of substrates. Beginning on Day 4, a significant difference in alignment is seen between cells seeded on random (0.08±0.01) and aligned (0.11 ±0.02) fibers. When cell alignment is analyzed at longer differentiation times through Day 14 and Day 28, cells differentiated on aligned and random fibers further diverge in their orientation indexes. By Day 14, cells differentiated on random fibers remain disorganized (S = 0.12±0.01), while cells differentiated on aligned fibers become significantly more aligned (0.17±0.01). This difference is further accentuated on Day 28, wherein cells differentiated on aligned (0.31±0.03) and random (0.17±0.02) fibers show an 80% difference in alignment (Fig. 2B).

**FIG. 2.**
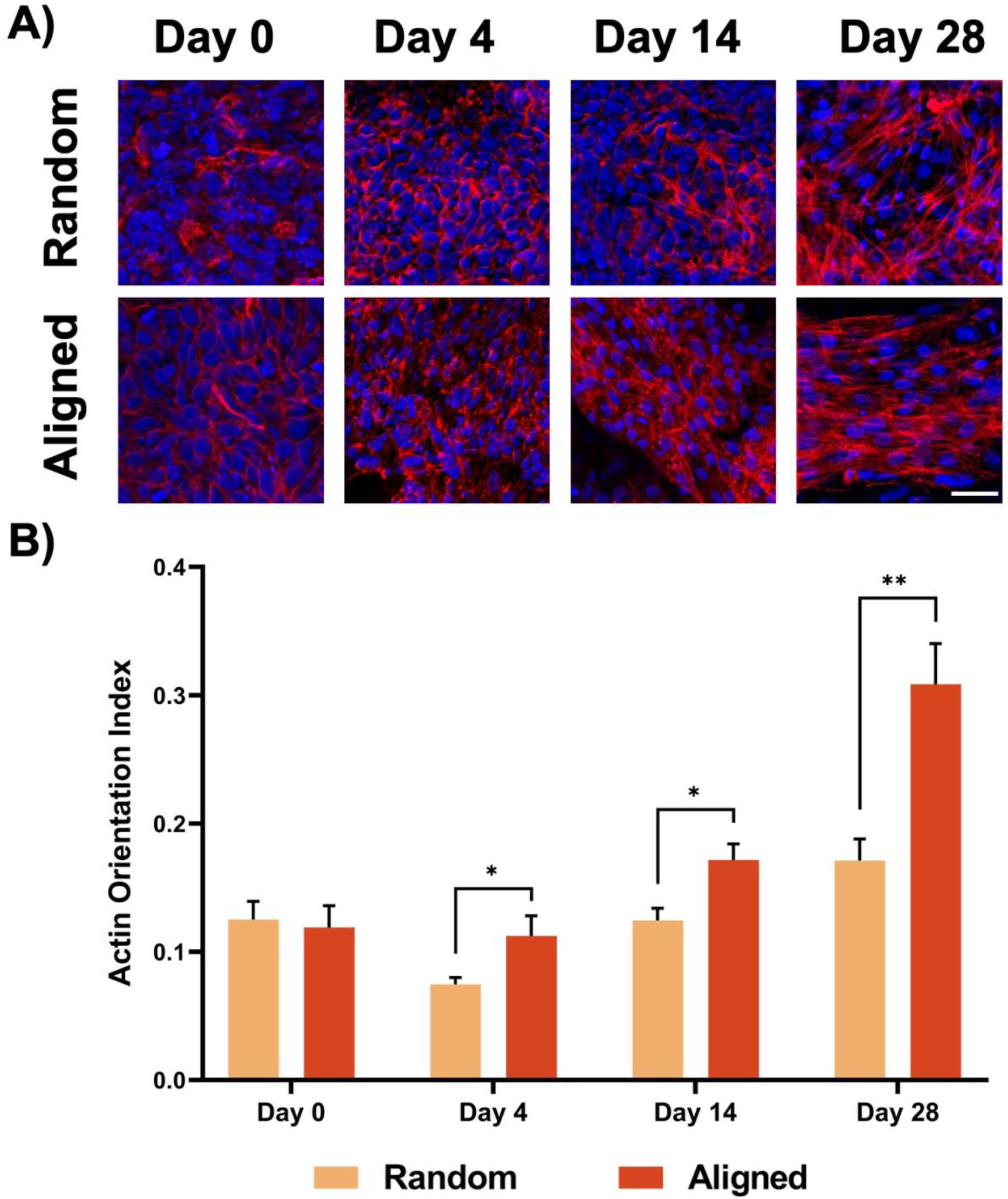
Aligned fibers engineer cell alignment during differentiation. **(A)** Nuclei (Draq4, blue) and actin filaments (Rhodamine phalloidin, red) of cells seeded on random or aligned fibers stained on day 0,4,14, and 28 of differentiation. Scale bar = 30μm. **(B)** Cell orientation index of hiPSCs seed on random and aligned fibers were calculated at day 0,4,14, and 28 of differentiation (n=30). *p<0.05,**p<0.01.

### Intracellular Calcium Transients

Next, we explored how cell alignment impacts cellular calcium handling. Following GCaMP imaging, we analyzed calcium release, flow, and reuptake rates at different time points during CM differentiation (Day 14 and Day 28). Synchronicity, a functional measurement of electro-coupling in the cardiac syncytium, was modulated by cell alignment in the short term (Fig. 3A). On Day 14, CMs differentiated on aligned fibers exhibited more synchronous calcium peaks (73.26±4.15) as compared to CMs differentiated on random fibers (Fig. 3C). At Day 28, the impact of fiber alignment disappears with CMs differentiated on both aligned and random fibers exhibiting similar levels of synchronicity (Fig. 3B). A similar trend is seen in beating frequency. On Day 14, CMs differentiated on aligned fibers had significantly lower BPM (77.8±1.65) than CMs differentiated on random fibers (93.15±2.26). By Day 28, BPM differences between alignment conditions are non-significantly different, with CMs differentiated on aligned fibers (61.88±1.34) only beating about 10% slower than CMs differentiated on random fibers (67.97±0.10) (Fig 3D). Alignment also drives the rate of calcium reuptake within CMs in the short term. The calcium cycle decay constant (τ), a measure of intracellular calcium reuptake, is seen to be significantly shorter at Day 14 for CMs differentiated on aligned fibers (283.70±11.32ms) as compared to CMs differentiated on random fibers (495.5±25.84ms). By Day 28, there is no significant difference between alignment conditions (Fig. 3E). Finally, the duration of the calcium cycle, or the full-width half-max of GCaMP traces, is driven by alignment at both time points. On Day 14, CMs differentiated on aligned fibers exhibit significantly wider FWHM compared to CMs differentiated on random fibers (222.70±4.24, 169±.4.24, respectively). This trend continues to Day 28, with the FWHM of CMs differentiated on aligned fibers (289.90±7.37) still significantly larger than CMs differentiated on random (224.50±4.74) fibers (Fig. 3F).

**FIG. 3.**
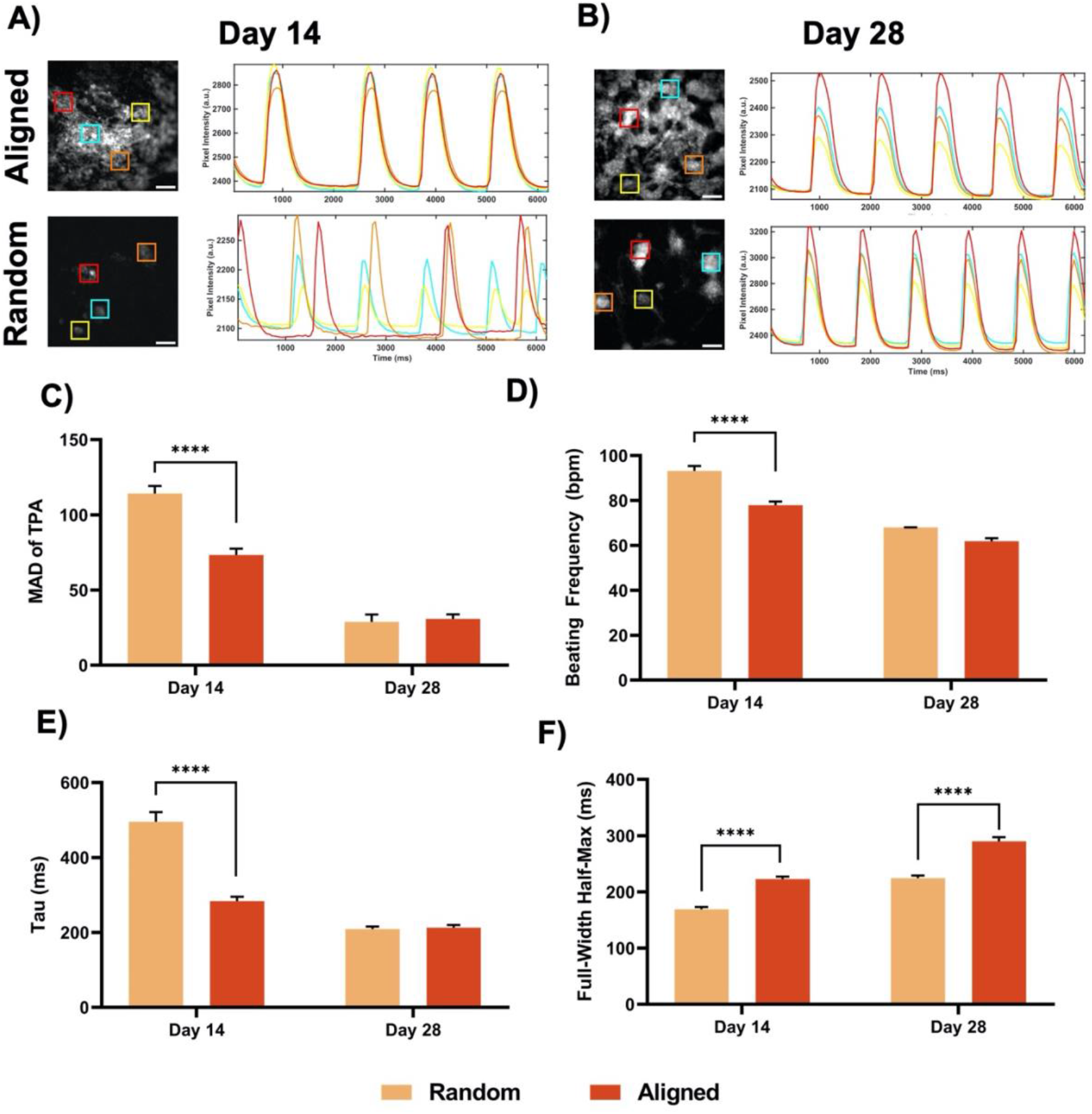
Intracellular calcium handling and synchronicity improve with engineered alignment. Representative calcium traces of GCaMP hiPSC-CMs differentiated on aligned and random fibers at day 14 **(A)** and day 28 **(B).** Scale bar = 200μm. **(C)** Synchronicity (n=90), **(D)** beating rate (n=96), **(E)** decay constant (n=96), and **(F)** calcium cycle duration (n=96) of day 14 and day 28 aligned and random hiPSC-CMs. ****p<0.0001. TPA-MAD, time of peak arrival median absolute deviation.

The above metrics show that functionally, CMs differentiated on aligned fibers exhibit improved calcium handling and coupling at earlier times compared to those on random fibers. These functions are supported by complex protein machinery. To identify the sources of these functional improvements, the expression of key cardiac and calcium handling genes was measured. On Day 14, no significant differences in calcium handling gene expression were seen between CMs differentiated on aligned and random fibers (Supplementary Fig. S3a). By Day 28, CMs differentiated on aligned fibers (2.38±0.47) show a significant upregulation in *CACNA1C* (encoding for the alpha1 subunit of voltage dependent calcium channel) compared to CMs differentiated on random fibers (0.81±0.15). (Supplementary Fig. S3b).

### Metabolism

CM metabolism was assessed using two optical techniques, one focusing on mitochondrial activity and the other focusing on mitochondrial size. Using ORR, the concentration of metabolic intermediates NADH and FAD+ were measured as indicators of the preferred path of ATP generation. The ORR was specified to CM activity via SIRPA, a cardiac-specific marker mask (Fig. 4A). The sensitivity of our analysis pipeline was confirmed with drug dosing of metabolic modulators for Day 14 differentiated CMs. Oligomycin A, an inhibitor of ATP synthase, is shown to significantly decrease the ORR of CMs differentiated on aligned fibers. Serial addition of FCCP to cells dosed with Oligomycin A increases ORR back to control levels (Supplementary Fig. S4b). This effect is similar to the trends seen in Oxygen Consumption Rate Seahorse assays.^21,26^

**FIG. 4.**
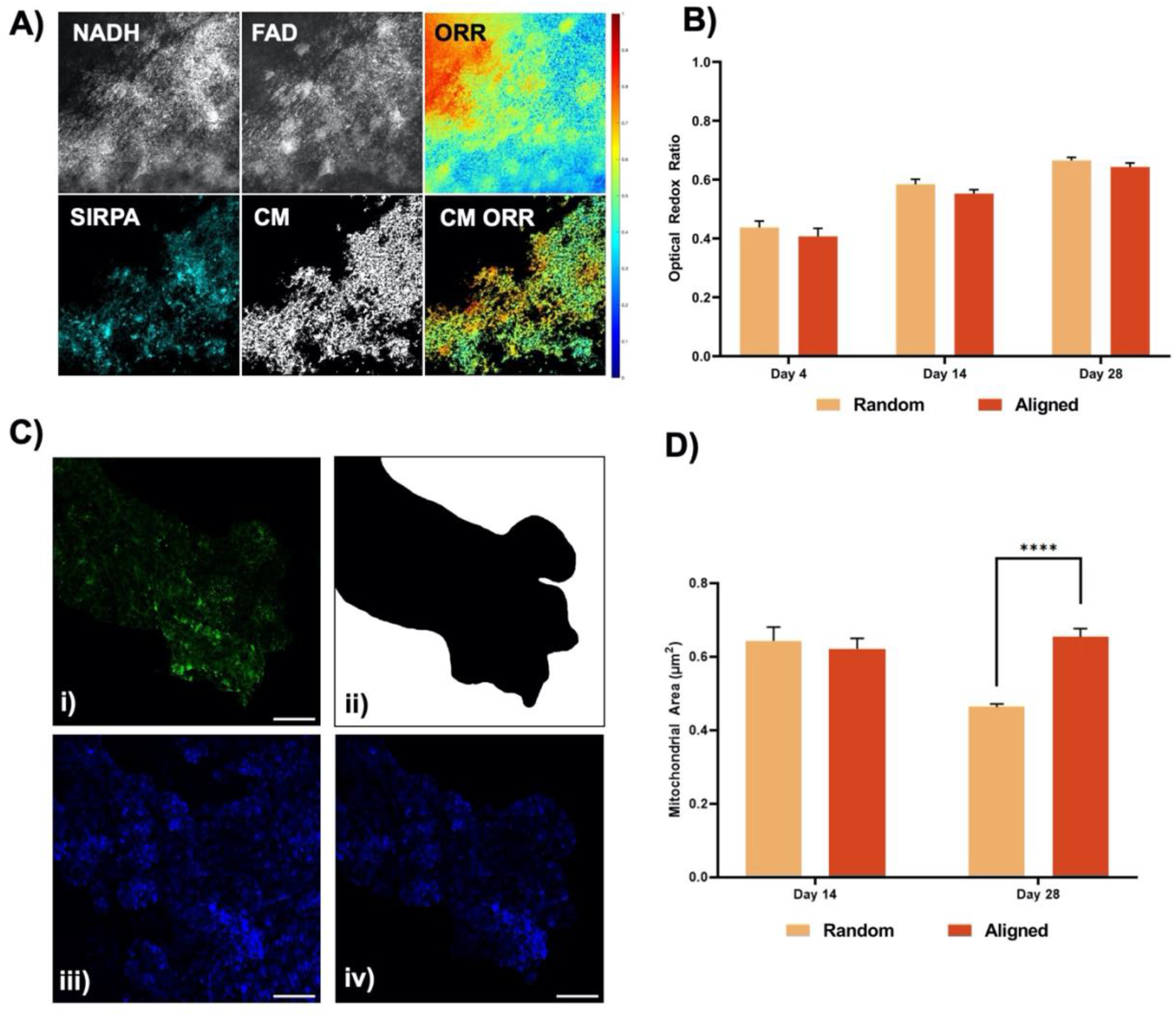
Metabolic analysis of hiPSC-CM throughout differentiation and development. **(A)** Representative images describing optical redox imaging and analysis needed to compute a cardiac specific optical redox ratio (ORR). Endogenous fluorescence of NADH and FAD are collected and used to calculate a non-specific ORR. Images of cardiac marker SIRPa are used to mask total ORR to CM-specific ORR. **(B)** Computed CM ORR for hiPSC-CMs differentiated on random and aligned fibers at days 4,14, and 28 (n=30). **(C)** Representative images showcasing analysis of hiPSC-CM mitochondria. Cardiac marker SIRPa images (i) are binarized (ii) and applied to fluorescent mitochondria images (iii) to generate cardiac specific mitochondrial images (iv). Scale bar = 50μm. **(D)** Average mitochondria size calculated for hiPSC-CM mitochondria on random and aligned fibers at days 14 and 28 (n=30). ****p<0.0001.

We next examined the impact of cell alignment on ORR at three time points: Day 4,14, and 28. Over time, all samples significantly increase in ORR, indicating a metabolic switch from glycolysis to oxidative phosphorylation. On Day 4, we did not observe significant differences in ORR of cells differentiated on random (0.44±0.02) or aligned (0.41±0.03) fibers. By Day 14, ORR for both samples increased by ~30% compared to Day 4 values; however, differences between CMs differentiated on random (0.58±0.02) or aligned (0.55±0.01) fibers prove to be insignificant. By day 28, ORR increases by 10-15% compared to day 14 and by 50% compared to day 4. Yet still, there are no significant differences between CMs differentiated on random (0.66±0.01) and aligned (0.64±0.01) fibers. (Fig. 4B).

Analysis of mitochondrial area reveals a long-term impact of cell alignment. On Day 14, the area of mitochondria within CMs differentiated on random fibers (0.64μm^2^) is not significantly different from those differentiated on aligned fibers (0.62μm^2^). On Day 28, CM differentiated on random fibers have a ~30% decrease in mitochondria (0.46μm^2^) area, while the mitochondria of CMs differentiated on aligned fibers (0.65μm^2^) remain close to Day 14 size (Fig. 4D).

Investigation of the expression of genes essential for glycolysis and fatty acid oxidation shows a short-term impact of alignment on metabolism. On Day 14, the glycolysis related gene *SLC4AC* and the FAO related gene *ACADL* are significantly upregulated in CMs differentiated on aligned (4.91±1.14) fibers compared to those differentiated on random fibers (2.31±0.23) (Supplementary Fig. S5a). By Day 28, these impacts disappear, with no significant difference in metabolic gene expression between CMs differentiated on random or aligned fibers (Supplementary Fig. S5b).

## Discussion

Herein, we investigate the effect of alignment on hiPSC-CM electrochemical coupling and metabolism. Previous work examined this impact by seeding terminally differentiated CMs onto aligned substrates; however, such an experimental design does not account for the natural CM alignment *during* maturation. As noted by many other studies, cell alignment can strongly affect mechano-sensing and cell-cell communication during differentiation.^27^ Our approach allows cells to sense substrate topology and alignment before and during differentiation. The feedback of this sensing is seen as early as Day 4 of differentiation as cells begin to align along the fiber direction. Additionally, other work uses purified CM-only samples to elucidate CM behavior and function, leveraging metabolic or cell sorting purification methods.^28^ Prior work has shown the existence and the importance of cross-talk between CMs and other cardiac tissue cell types like fibroblasts and endothelial cells.^29^,^30^ Our approach preserves a heterogeneous cell population after differentiation, enabling cellular cross-talk and allowing us to measure the impacts of this environment on CMs, using canonical CM markers CTnT and SIRPa to identify CMs for cell-type-specific analysis.

Using engineered cell alignment during (not after) hiPSC-CM differentiation and maturation we can compare electrochemical coupling and metabolic effects of alignment. Our analysis reveals longitudinal impacts of alignment on CM calcium handling and metabolism. Adult, healthy CMs show structured alignment of sarcoplasmic reticulum and gap junctions that engender efficient calcium handling and signal propagation.^3^ We hypothesized that engineering alignment in hiPSC-CMs would improve calcium handling. Our analysis measured metrics of both intracellular calcium handling and intercellular calcium flow. We measured beating frequency (to estimates the leakiness of potassium and calcium ion channels), in hiPSC-CMs the FWHM of calcium cycle (to measure the duration of elevated intracellular calcium levels and the function of SERCA pump proteins), and Tau, (the decay constant of calcium reuptake by the sarcoplasmic reticulum). Finally, we quantified beating synchronicity to assess intercellular hiPSC-CM communication. In general, hiPSC-CM maturity is associated with lower BPM, longer FWHM, smaller decay constants, and higher levels of synchronicity.^3^,^26^ Our data show that alignment improves calcium handling maturity in the short term in all four metrics compared to CMs differentiated on random fibers. However, by Day 28, CMs differentiated on random fibers have the same performance as CMs differentiated on aligned fibers in all metrics except calcium cycle duration (Fig. 4). These results align with our previous work with mouse embryonic stem cell-derived CMs.^24^ Similarly, the positive impact of alignment on human CM calcium handling is well supported in the field.^31^ Notably, our data on calcium cycle duration supports the connection between cellular disorganization and SERCa2a dysfunction in cardiomyopathies.^32^ Several factors may contribute to the long-term decoupling of alignment and hiPSCs-CM calcium handling we see in our study. First, cell confluence may drive long-term EC coupling and calcium handling. Multiple studies connect CM contraction to mechanical forces exerted by neighboring cells.^33,34^ Nitsan et al. showed that mechanical stimulation, mimicking the contractions of neighboring cells, can pace CM contraction. Thus, at later time points, high cell confluence may drive the synchronicity of random CMs more strongly than their alignment. Next, the presence of non-CMs may also contribute to improved calcium handling in the long term, for which alignment may have an impact. We generate a heterogeneous cell population as we do not include a purification step during CM differentiation. We observed, via immunostaining, the presence of fibroblasts in our culture after differentiation, with lower amounts of endothelial and smooth muscle cells. Fibroblasts, cocultured with CMs, are well known to improve CM electrophysiology.^35,36^ Compared to hiPSC-CMs, fibroblasts experience dynamic growth over time, changing with development^37^ and disease.^38^ Future work should evaluate how alignment impacts the growth kinetics of fibroblasts to determine if population changes are responsible for long-term equalization of calcium handling between random and aligned tissues.

We next studied how cell alignment may impact CM metabolism. Mature CMs feature structured intracellular energetic units (IEUs) that preferentially use oxidative phosphorylation for ATP synthesis. These units, composed of tightly packed and aligned sarcomeres, t-tubules, sarcoplasmic reticulum, and mitochondria, ensure the efficient transport of ATP within CMs.^39^ It has been shown that IEUs effectively integrate mechanical stimuli into improved function, but studies have tested how tissue alignment factors into this process.^15^ Given the strong structure-function relationship in healthy cardiac tissue, we hypothesized that CM alignment would improve mitochondrial function and accelerate the metabolic switch in CM development. Our assessment of CM metabolism allowed us to track mitochondrial size and the production of metabolic intermediates optically. We expected CMs differentiated on aligned fibers to be more metabolically mature with larger mitochondrial area and higher ORR than CMs differentiated in random fibers. Our hypothesis was correct regarding the mitochondrial area, but ORR measurements showed no effect of cell alignment on CM metabolism. Yet, a metabolic switch from glycolysis to oxidative phosphorylation is seen in both samples over time. Future studies will focus on utilizing ORR to study how alignment in conjunction with fatty acid supplementations can impact CM metabolism.

Changes in the average mitochondrial area of random CMs lend support to the interrelation of cytoskeletal structure and mitochondrial morphology. This relation is clear in cardiac disease states; various cardiomyopathies exhibit myofibril disorganization concurrent to mitochondrial fragmentation.^40–43^ Interestingly, we see a disconnect between mitochondrial morphology and electrochemical coupling for random CMs on Day 28. Future studies will include drug modulation of mitochondria to further elucidate the relationship between electrochemical coupling and mitochondrial organization.

In summary, our studies show that differentiating hiPSC-CMs on aligned fibers accelerates initial electrochemical coupling and matures calcium handling. In addition, we find that cell alignment (in a heterogeneous population) does not impact CM metabolism switch but may affect mitochondrial fission and fusion rates. While ORR has been mainly used for cancer cell studies until now, here we demonstrate the utility of using ORR for measuring metabolic changes during CM differentiation, which is also critical for CM maturation. We see these aligned tissues as ripe platforms upon which other known maturation inputs may be deployed to trigger multidimensional hiPSC-CM maturation.

## Acknowledgements

We gratefully acknowledge the support of the National Science foundation (2002652, awarded to J.Z.), the Welch Foundation (F-2008-20190330, awarded to S.H.P.) and the Human Frontiers in Science Program (RGP0045/2018, awarded to S.H.P.). The authors thank Dr. Elizabeth Cosgriff-Hernandez for use of the Phenom Pro scanning electron microscope, Ali Mohsin for help with SEM image processing and fiber alignment quantification, and Dr. Nima Momtahan for aid in hiPSC-CM differentiation.

## Author Contributions

**Nikhith G. Kalkunte:** Conceptualization, Methodology, Software, Writing – Original Draft. **Talia E Delabmbre:** Investigation, Validation. **Sogu Sohn**: Validation, Writing – Original Draft. **Madison Pickett:** Validation, Resources. **Sapun Parekh:** Conceptualization, Writing - Review & Editing. **Janet Zoldan:** Conceptualization, Writing - Review & Editing, Supervision

