## Supplementary figures and images for "Engineering Alignment has Mixed Effects on hiPSC-CM Maturation"

### Supplementary Figure 1

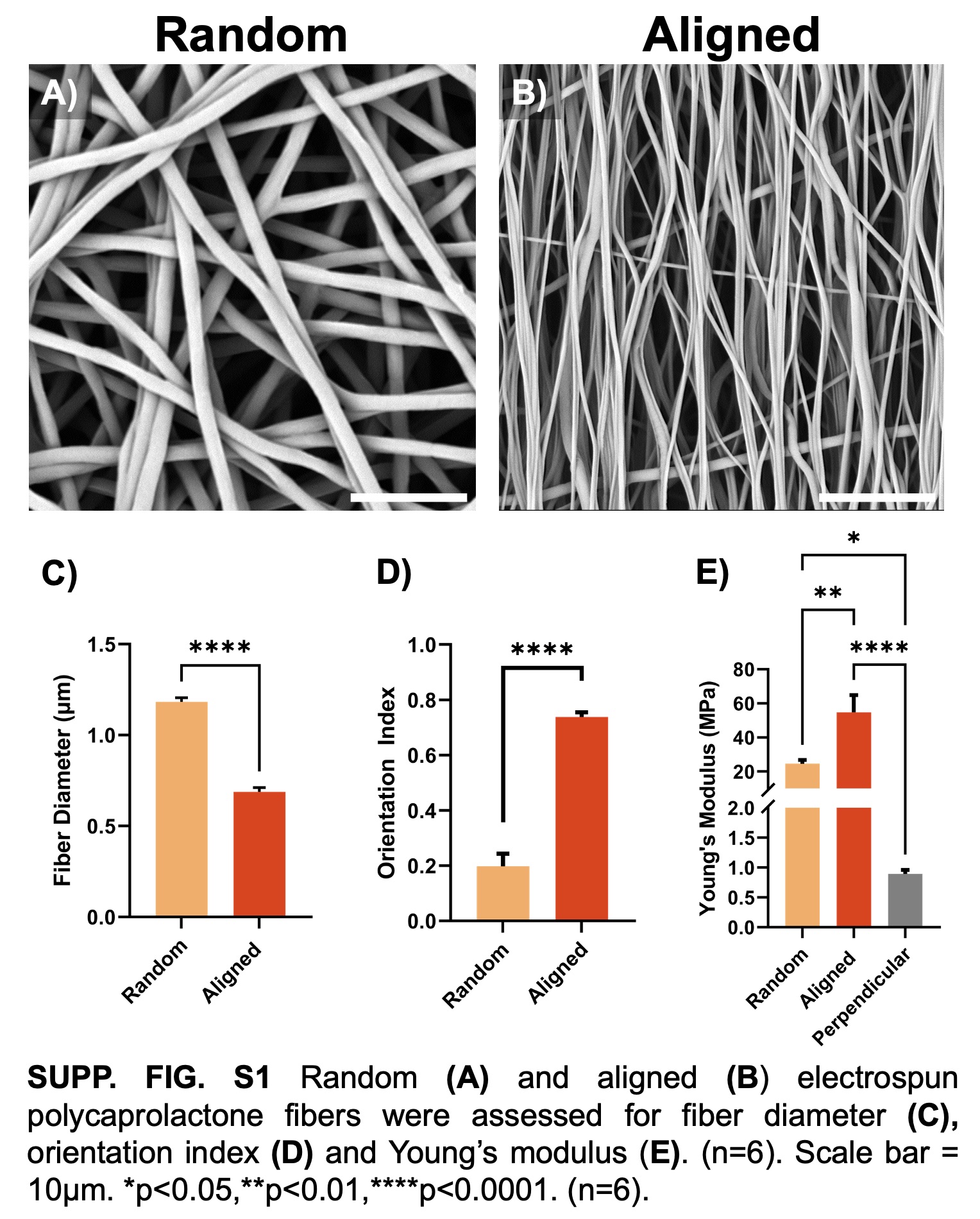

### Supplementary Figure 2

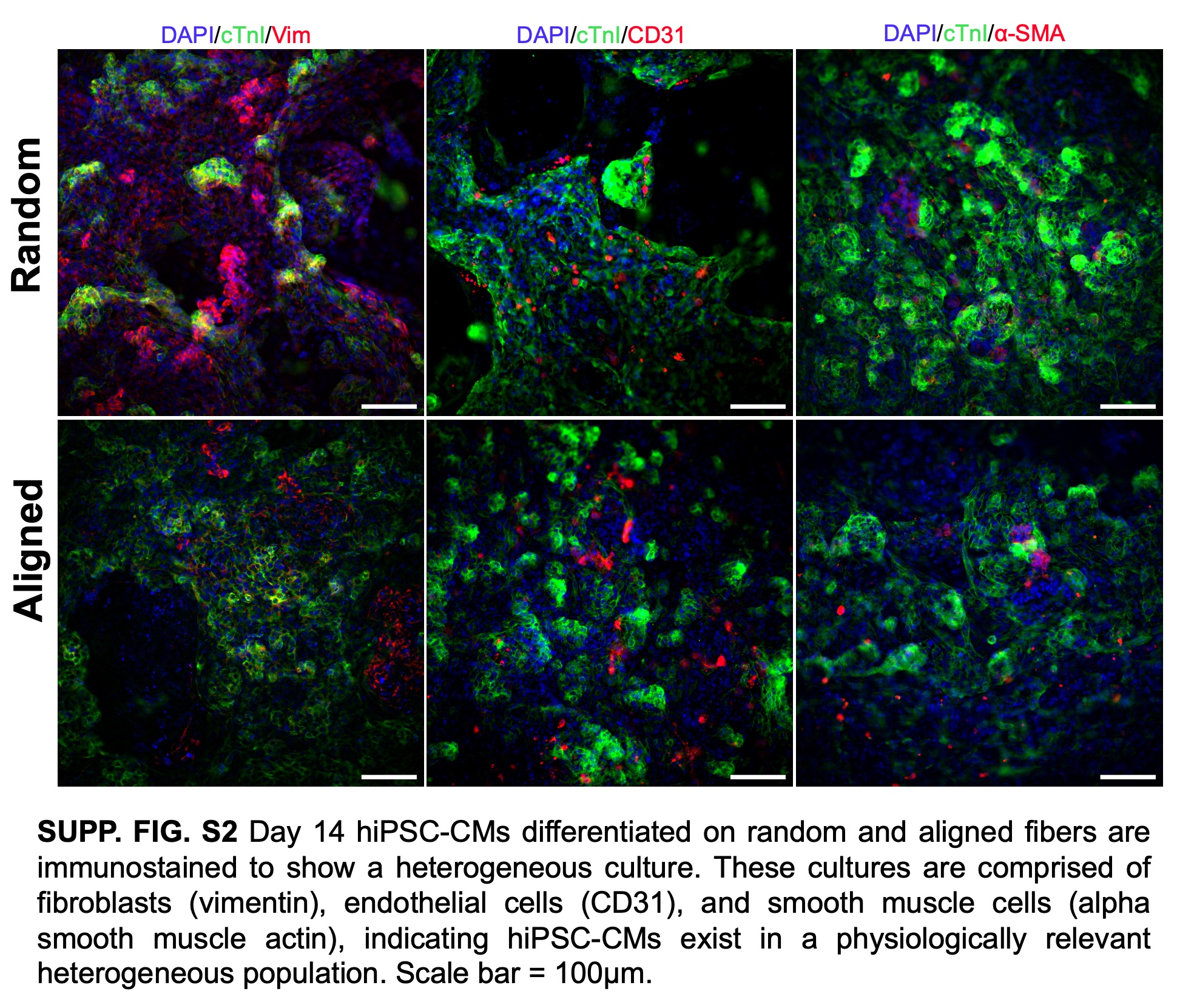

### Supplementary Figure 3

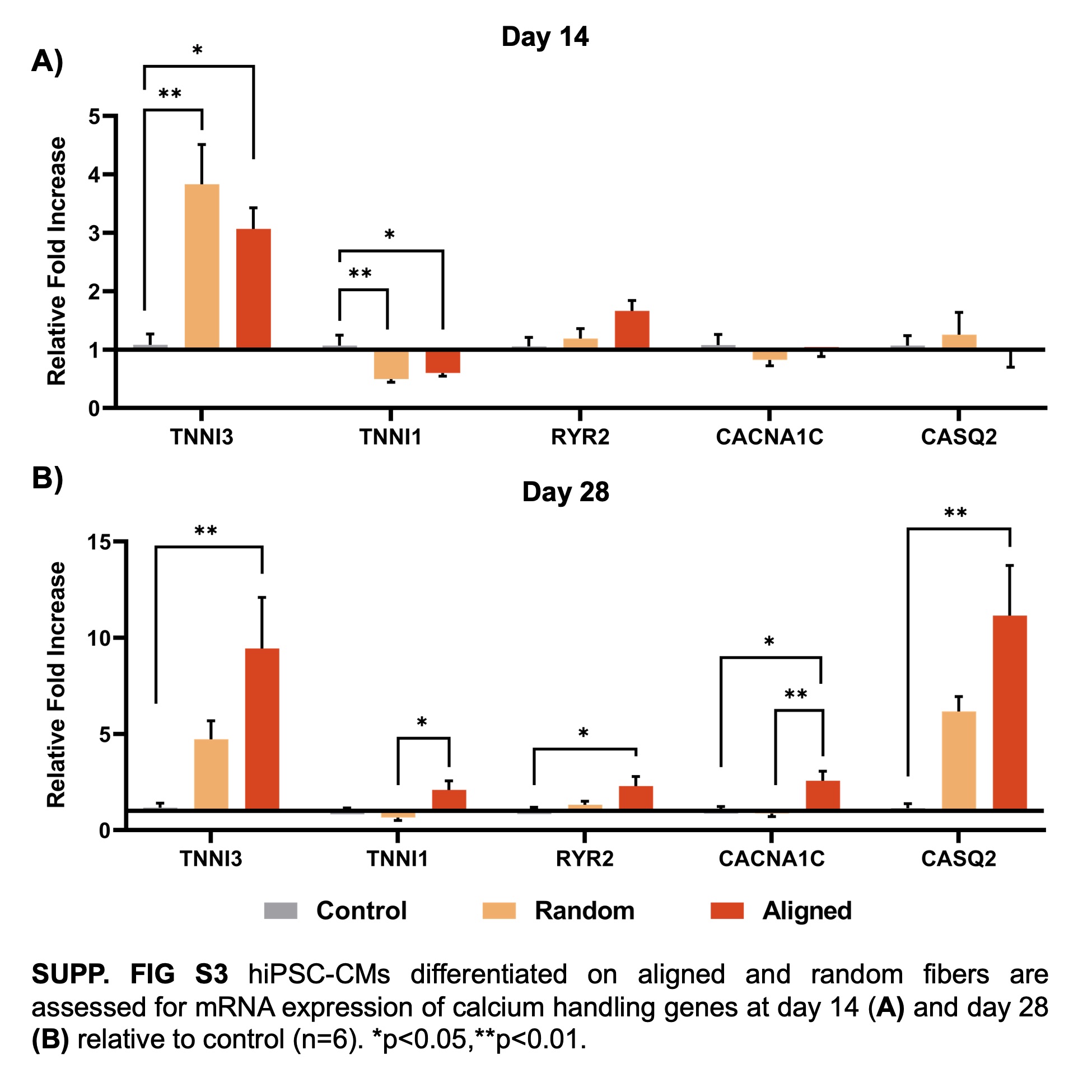

### Supplementary Figure 4

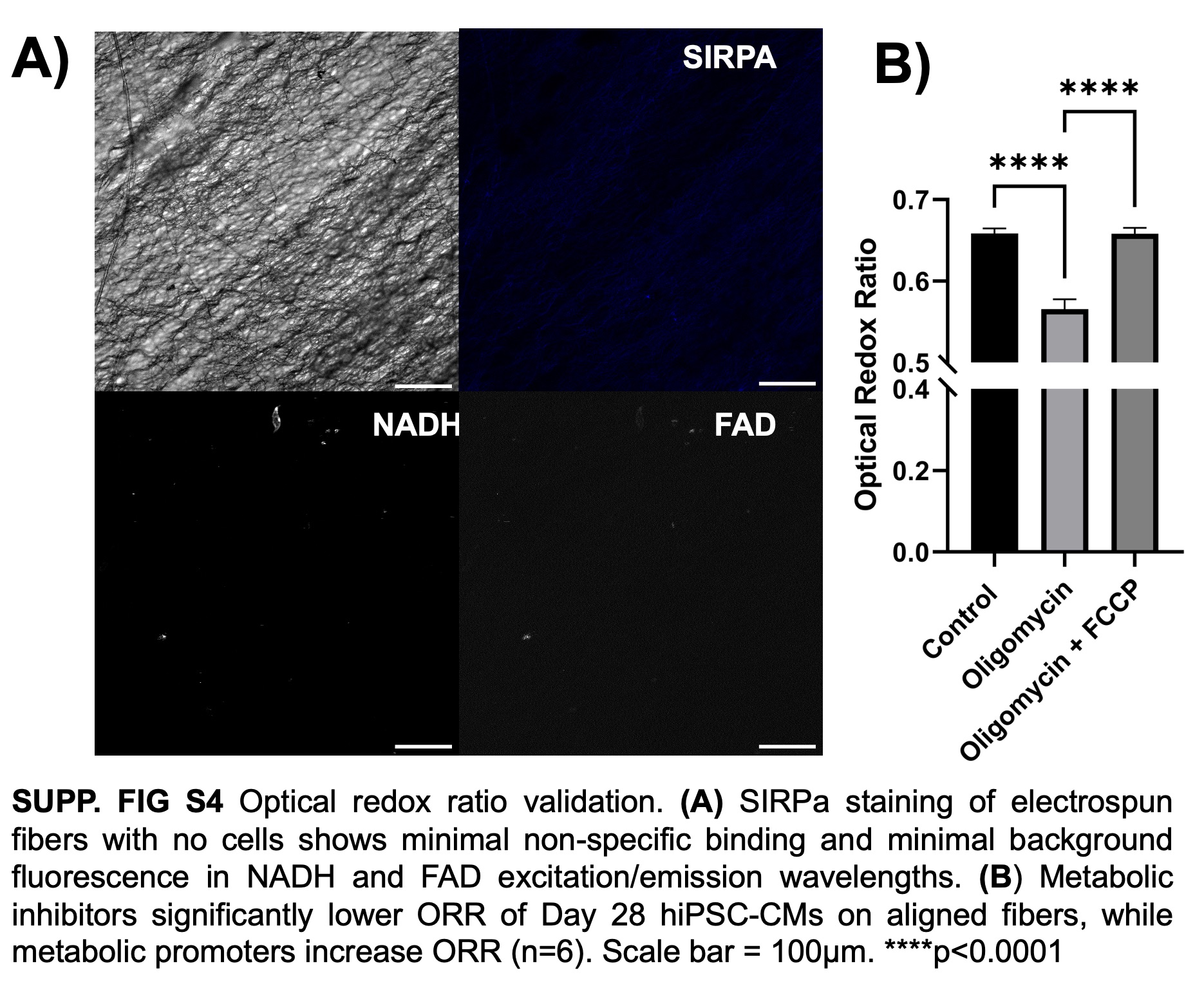

### Supplementary Figure 5

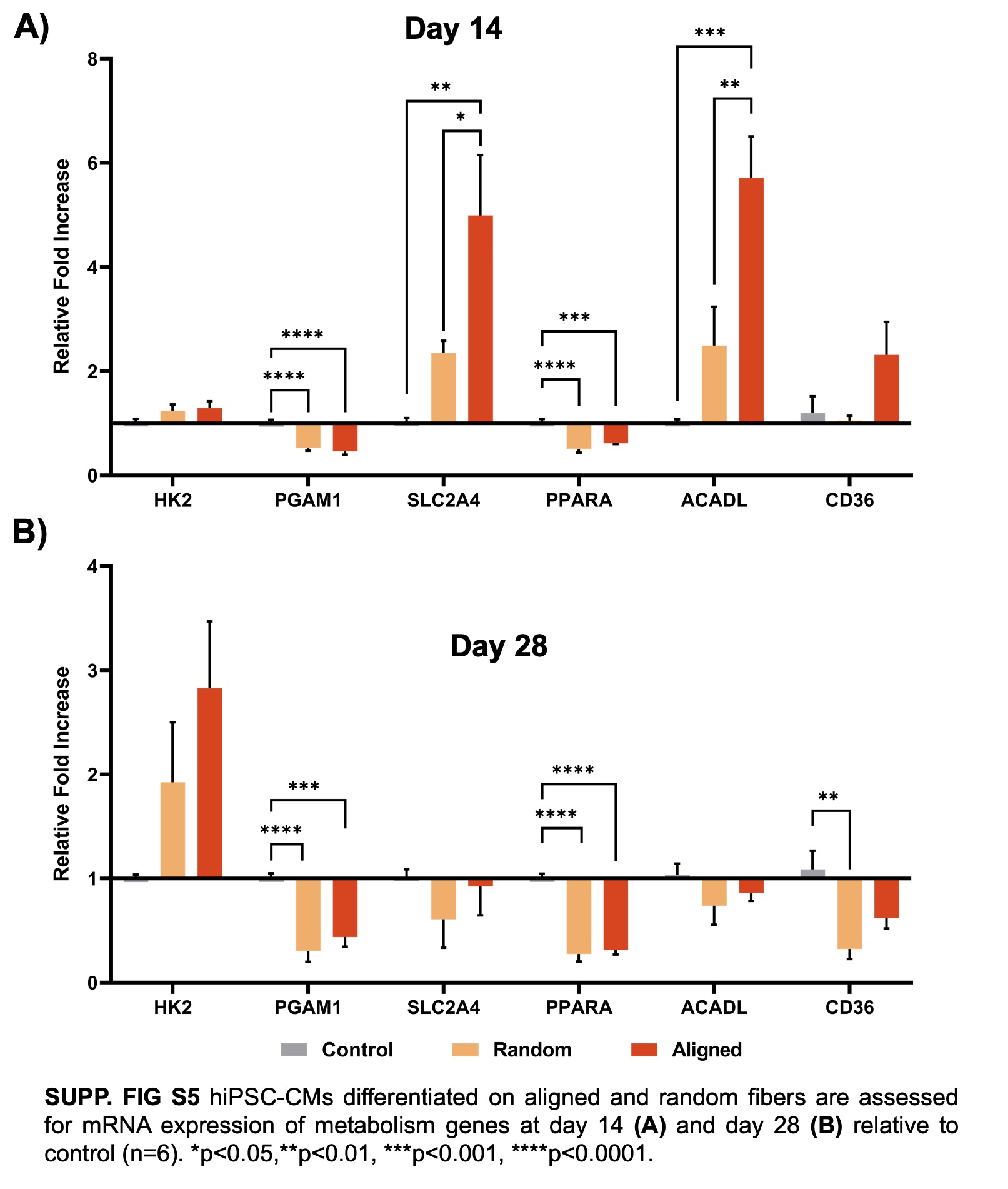

### Supplementary Table 1

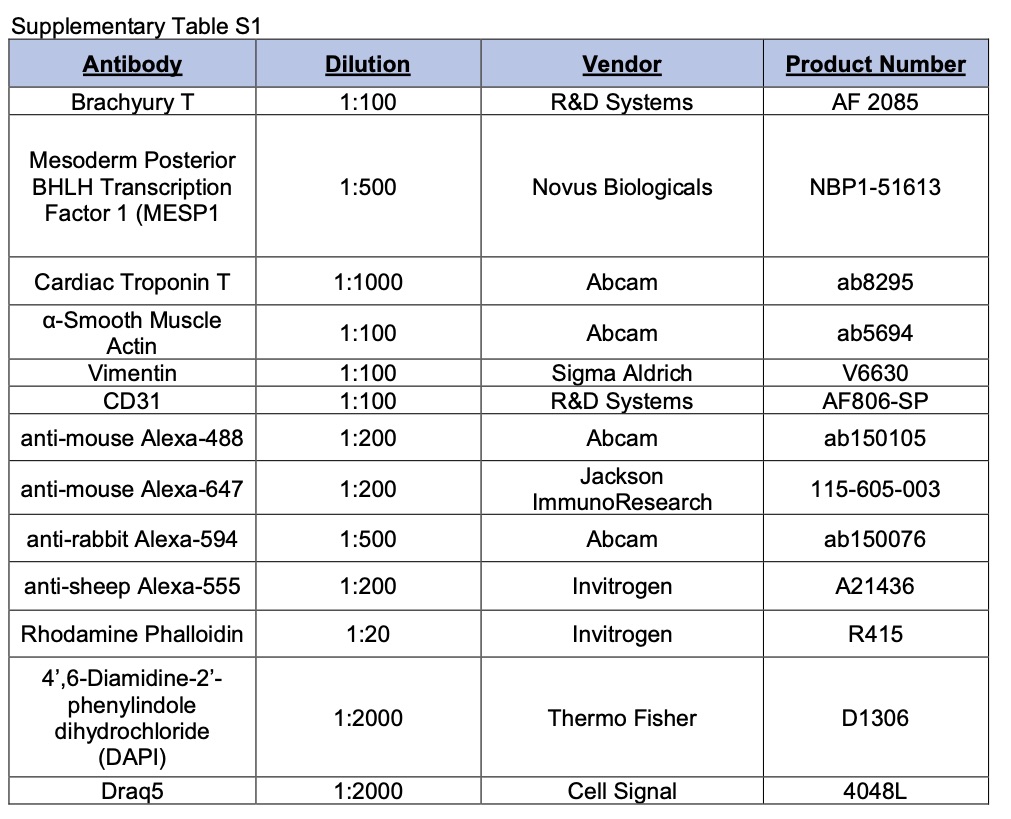

### Supplementary Table 2

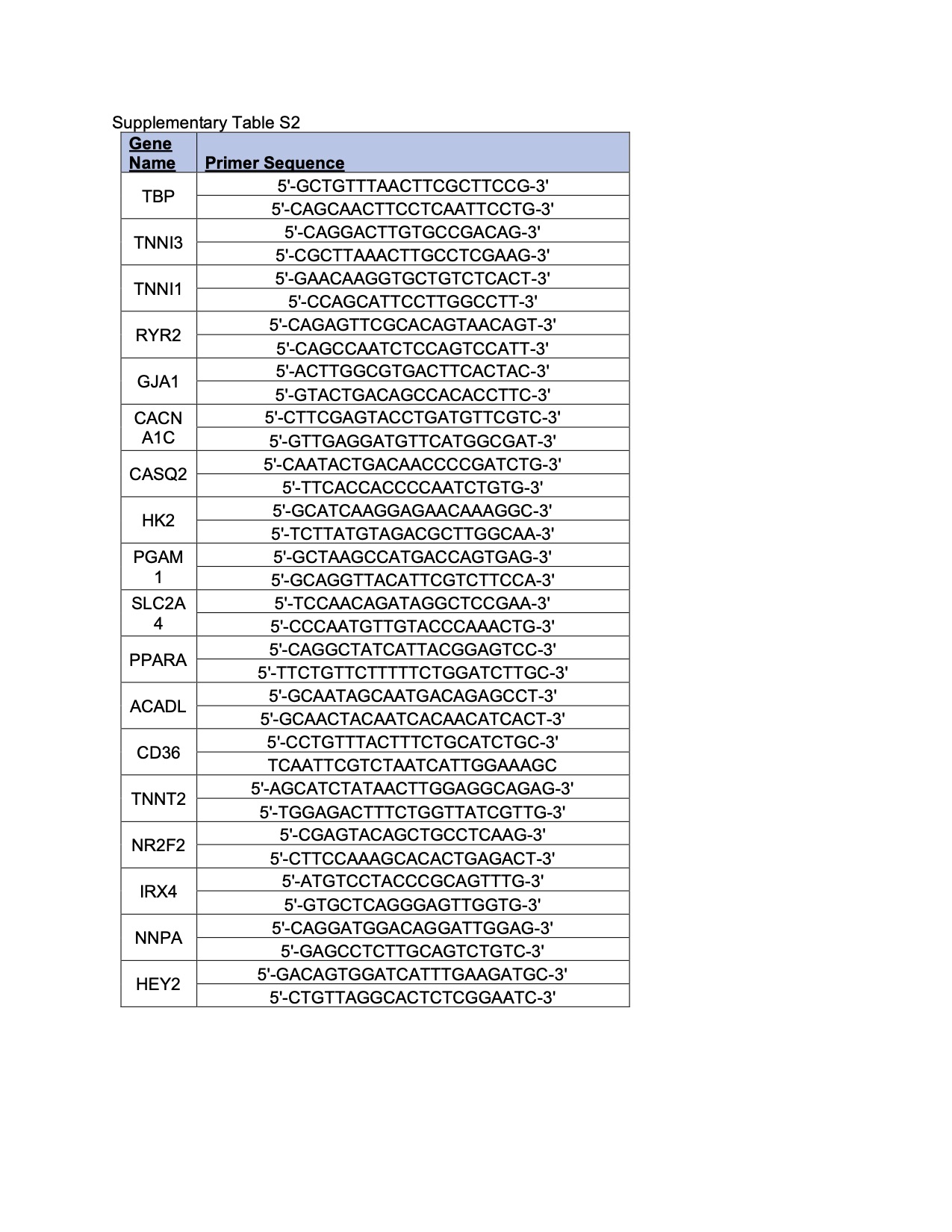
